# Illustrating Potential Effects of Alternate Control Populations on Real-World Evidence-based Statistical Analyses

**DOI:** 10.1101/2021.02.21.432172

**Authors:** Yidi Huang, William Yuan, Isaac S. Kohane, Brett K. Beaulieu-Jones

## Abstract

Case-control study designs are commonly used in retrospective analyses of Real-World Evidence (RWE). Due to the increasingly wide availability of RWE, it can be difficult to determine whether findings are robust or the result of testing multiple hypotheses. We investigate the potential effects of modifying cohort definitions in a case-control association study between depression and Type 2 Diabetes Mellitus (T2D). We found that small permutations to the criteria used to define the control population result in significant shifts in both the demographic structure of the identified cohort as well as the odds ratio of association. These differences remain present when testing against age and sex-matched controls. We believe this offers strong support for the need for robust guidelines, best practices and regulations around the use of observational RWE for clinical or regulatory decision making.

## Introduction

The FDA has shown a strong interest in the utilization of Real-World Evidence (RWE) to enhance or replace aspects of the regulatory process [1,2]. Recently the FDA increased their participation in a partnership in RWE for oncology [3] and the pace of accelerated approvals has increased substantially from 2012 [4,5]. These actions have been met with mixed reactions [6], especially regarding attempts to replace traditional randomized controlled trials (RCTs) with RWE-based comparative effectiveness analyses [7,8]. The important aspects of the utilization of RWE in the regulatory process we consider in this paper are: 1.) lack of pre-registration, 2.) protection against intentional or unintentional multiple testing, and 3.) the potential for financial incentives to drive the strategic selection of a cohort given prior testing of retrospective data.

The digitization of medical records and administrative data have made research using RWE increasingly prevalent, and RWE has the potential to be an incredible resource to the research community [9,10]. RWE has already enabled the study of patient-level health outcomes at an unprecedented scale, with innovative study designs that address questions such as 1.) genetic heritability of different phenotypes with large scale twin studies [11], and 2.) prescribing patterns in the opioid epidemic [12].

Due to the fact that RWE is readily available, with numerous datasets that can be purchased from commercial providers (e.g. IBM Marketscan [13], Optum [14], Premier Healthcare Database [15]), it is not possible to enforce traditional pre-registration that typically accompanies active enrollment comparative effectiveness studies (e.g. RCTs). Because of this lack of pre-registration, it is more difficult and potentially impossible to determine if multiple groups have tested the same hypothesis especially given the publication bias towards positive results [16]. When the inability to detect multiple testing is combined with the financial incentives inherent to the regulatory process for both new therapies and post-approval surveillance, there exists a possibility for bad actors to exploit the ability to intentionally multiple test or “p-hack” [17]. Evidence to the potential for actions of this nature can be seen in recent events such as the data manipulation that occurred during the approval of Zolgensma [18,19]. It is important to note that current RCTs are not immune to selection bias as demonstrated by the fact that they commonly include younger [20] healthier participants [21]. This leads to generalizability concerns for RCTs but at least in this case, due to randomization, there is no ability to pre-select the case or control groups.

In this work we demonstrate the ability to profoundly affect the odds ratio for an association between depression and Type II Diabetes Mellitus (T2D) by making small, justifiable alterations to the control definition using published T2D phenotyping algorithms from eMERGE [22]. These algorithms were originally designed to overcome the challenges in identifying patient cohorts in EHR [23]. We hypothesized that the requirement for a glucose test in the eMERGE may select for controls that are less healthy on average than the overall potential control population. We considered a previously published association between depression and T2D and evaluate how the association changes with small permutations to the control population for T2D. Comorbidity between T2D and depression is well documented [24–26]. The causal relationship between these diseases is best characterized as complex, and evidence exists to support both that depression elevates risk of T2D and vice versa, as well as the hypothesis that both diseases share common etiology [25]. This experiment demonstrates a need to show that findings derived from RWE are robust to small permutations in the included population.

Existing literature on case-control methodology has focused on confounding due to lack of randomization as well as biases in study design [6–8,27]. Confounding due to unmeasured variables is a major concern, and makes it difficult to determine causal relationships from observational data [7]. Other pitfalls include the nonrandom assignment of exposures [6] and the nonrandom selection of participants [27]. Control selection has been identified as a crucial component of case-control study design, although best-practice recommendations are centered around matching techniques to control for confounding effects [28,29]. Multiplicity is discussed in the context of multiple hypothesis testing in genetic association studies [30], but we were unable to find existing work studying permutations in study design. Previous work has shown that the modeled effect size of mortality risk factors can be profoundly sensitive to model selection [31], suggesting to us that association results may also be sensitive to permutation in study design.

## Methods

We performed all analyses using a large de-identified administrative claims dataset including more than 75 million individuals for the timeframe from January 1, 2008, through June 30, 2017. This database does not include any race or ethnicity data and its usage has been deemed to be deidentified non-human subjects research by the Harvard Medical School Institutional Review Board, therefore waiving the requirement for approval. The database includes member age, biological sex, and enrollment data, records of all covered diagnoses and procedures as well as medication and laboratory results for a substantial subset of the covered population.

An overview of the study design is shown in Figure 1. To match cohorts on coverage status, members were required to have at least four years of continuous enrollment to qualify, and only the first four years were considered when defining phenotypes. Case and control populations were generated by applying various definitions (Figure 2) to the first four years of claims. Case and control cohorts were sampled from populations of various sample sizes (n=1000, 2000, 5000, 10000) to test for association between depression and T2D status. Association testing was performed using Fisher’s exact test to calculate the odds ratio (OR) and associated p-value. Each test was resampled 200 times using the bootstrap method to obtain a sampling distribution for the OR. Tests were performed with and without age and sex matching and the results compared.

**Figure 1.**
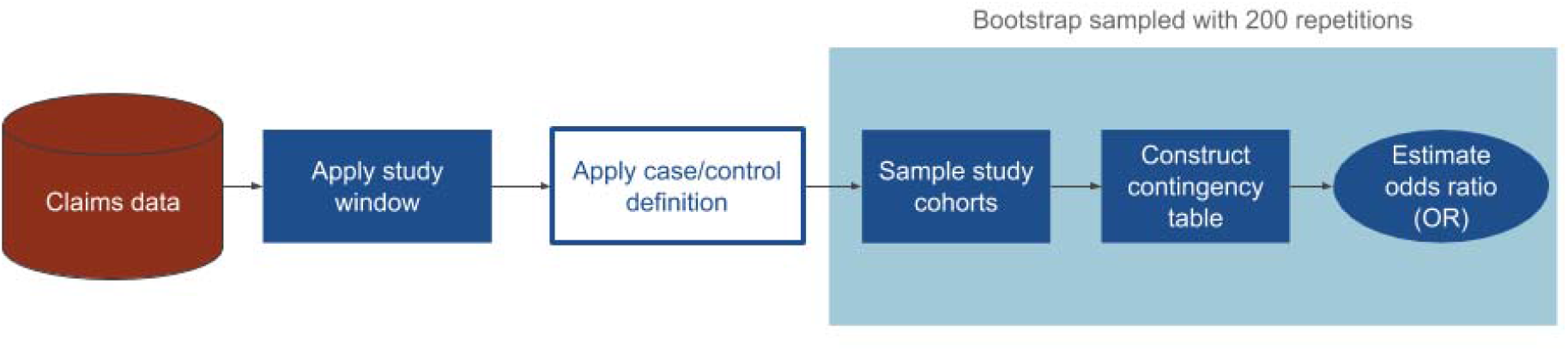
Overview of study design to evaluate the effect of alternate control groups on the association between depression and Type 2 Diabetes Mellitus.

**Figure 2.**
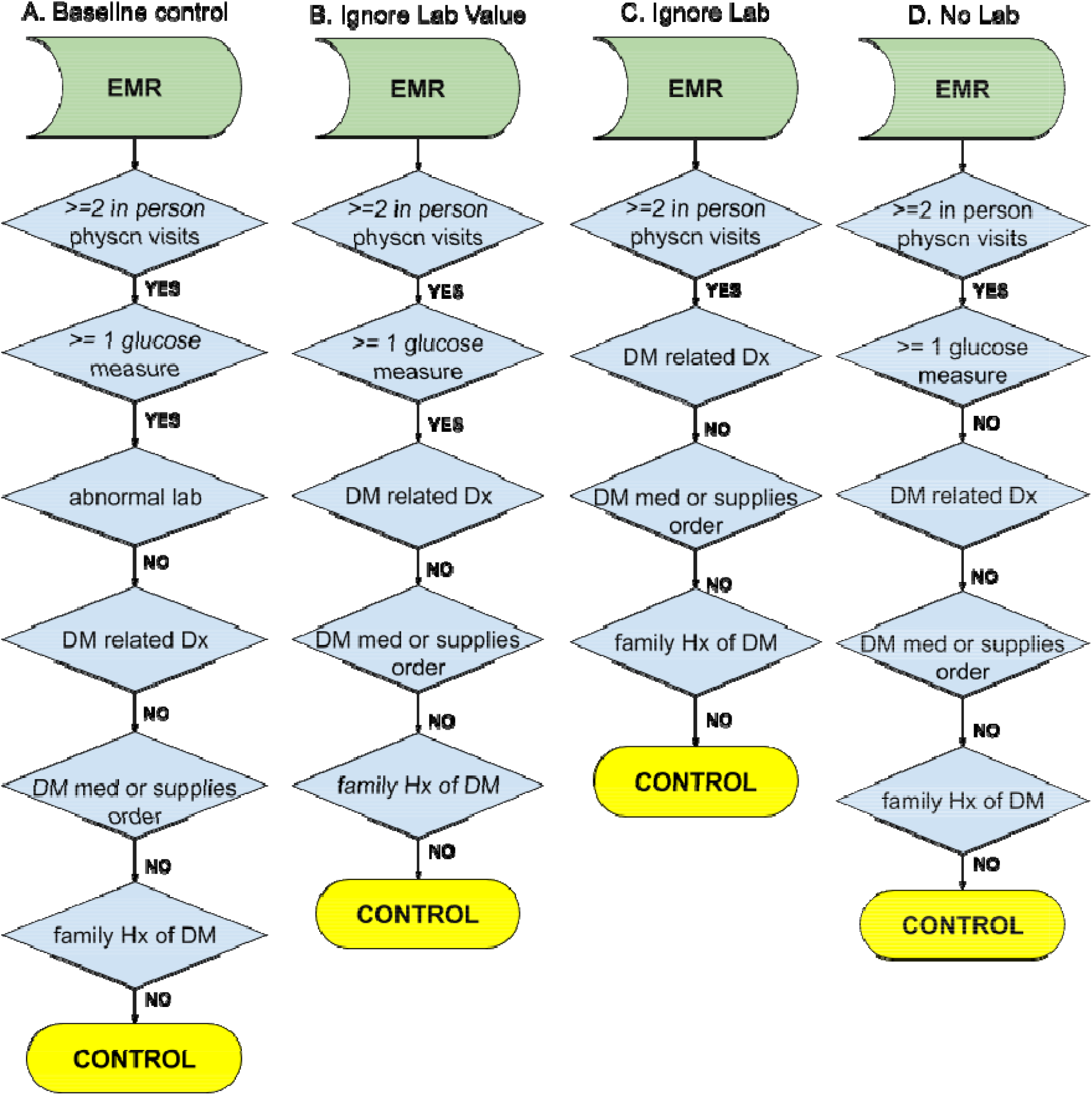
Control algorithms for Type 2 Diabetes Mellitus. A.) Baseline controls as defined by eMERGE. B.) Controls where the glucose lab value is ignored. C.) Controls where whether the member has had a glucose lab or not is ignored. D.) Controls where the member has not had a glucose test.

T2D Case status was determined using an adaptation of the eMERGE T2D phenotyping algorithms [22] for claims data. Each distinct claim was considered a separate visit for the determination of visit count. Diagnoses and medications were determined from medical and pharmacy claims, respectively. Ingredient-level RxNorm codes were mapped to NDC codes using the RxNorm API [32]. In the absence of clinical notes or structured questionnaires, family history was determined using ICD code V18.0 in medical claims. This is a limitation of using RWE to consider comprehensive medical histories.

Multiple control groups were defined based on the eMERGE T2D control algorithm with variations on the lab testing requirement (Figure 2). The eMERGE algorithm considers tests for fasting glucose (LOINC 1558-6), random glucose (2339-0, 2345-7), and hemoglobin a1c (4548-4, 17856-6, 4549-2, 17855-8) with defined thresholds for abnormal values. We include these LOINC codes when considering lab values. Within the claims data, all lab orders are available but only ∼20% of lab values are passed back to the private insurer. When checking to see if a lab test is ordered, we add procedural (CPT 82947, 80047, 80048, 80053, 80069, 83036) codes to determine whether a test was ordered. The groups are defined as follows:

1. eMERGE T2D control (labeled “Baseline control”)
2. eMERGE T2D control still requiring lab test presence but without requirement on lab test result (labeled “Ignore Lab Value”)
3. eMERGE T2D control without requirement on lab test presence (labeled “Ignore Lab”)
4. eMERGE T2D control requiring no lab test (labeled “No Lab”)

Exposure status for depression was defined using the CCS rollup for depressive disorders (single-level CCS diagnosis category 6572). Members with a qualifying enrollment period without a depression diagnosis were defined to be non-exposed for the purposes of this association with the acknowledgment that this does not rule a diagnosis prior to enrollment but indicates that there is not active care being provided for depression.

All analyses were performed using queries in Microsoft ® SQL Server 2017 and Python™. Statistical calculations were performed using Numpy and Scipy. Visualizations were created using Matplotlib and Seaborn. All source code is available in archival form on Zenodo [33] and on Github as a Jupyter notebook https://github.com/brettbj/association-robustness.

## Results

Table 1 summarizes the demographic characteristics of the different groups. Fact count is determined as the total number of ICD codes recorded per patient per year in the qualifying four-year window. Cases are on average older and have higher fact count than controls. Furthermore, controls who had lab testing are older still and have higher fact count than those who did not have lab testing.

**Table 1.** Demographic statistics under different control definitions

|  | Case Population | Baseline control | Ignore Lab Value | Ignore Lab | No Lab |
| --- | --- | --- | --- | --- | --- |
| Members | 289,113 | 2,288,032 | 2,547,932 | 8,062,907 | 3,397,361 |
| % Male | 52.52 | 43.11 | 43.75 | 48.09 | 52.01 |
| % Female | 47.48 | 56.89 | 56.24 | 51.91 | 47.99 |
| Age | 65.11 (12.19) | 49.96 (16.72) | 50.89 (16.89) | 43.34 (21.86) | 32.73 (20.86) |
| % Members with Depression | 16.38 | 13.83 | 14.07 | 9.96 | 4.19 |
| Total Facts | 50.06 | 25.27 (28.09) | 26.67 (30.62) | 20.40 | 11.03 (14.16) |
| Per Year | (50.31) |  |  | (26.84) |  |

Figure 3 shows that members receive more glucose testing as they age (per member per year). This indicates that requiring a glucose test may cause the control population to be older on average. Indeed, the control population with glucose testing is on average 18 years older than the control population without. This is one indication that requiring glucose testing as part of the eMERGE control algorithm may select for an older, less healthy population than the entire potential control population.

**Figure 3.**
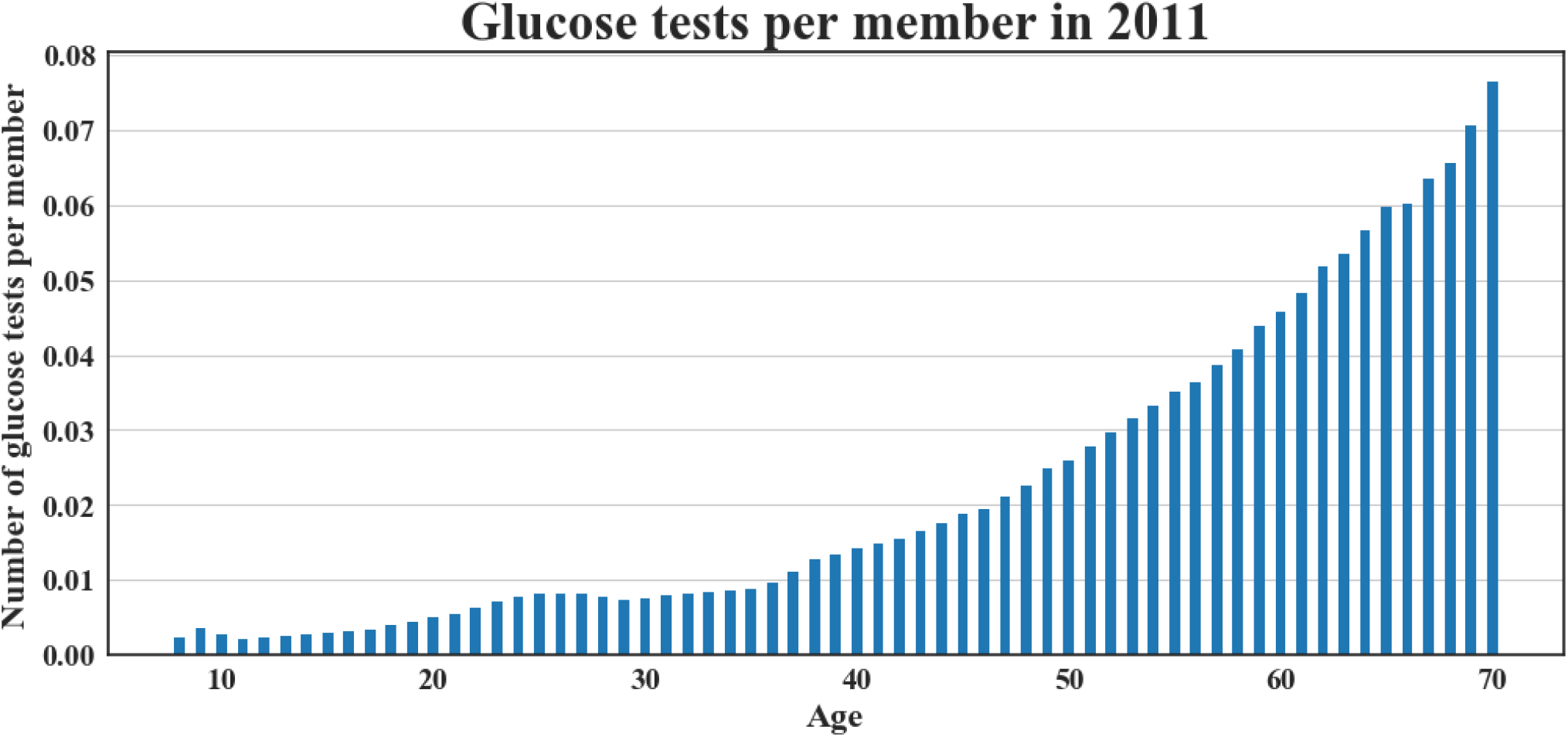
Number of glucose tests per member during 2011 by age.

Adjusting the glucose lab requirement in the control definition results in different age distributions (Figure 4). Matching on age and sex corrects for these differences but the percentage of cases with depression diagnoses and the total number of diagnoses, or facts, is higher in the case population (Table 2).

**Table 2.** Case and Control Summary Statistics after Matching.

|  | Case Population | No Labs Matched |
| --- | --- | --- |
| Members | 285,112 | 285,112 |
| % Male | 51.85 | 51.85 |
| % Female | 48.14 | 48.14 |
| Age | 65.01 (12.25) | 65.01 (12.25) |
| % Members with Depression | 16.45 | 6.60 |
| Total Facts Per Year | 50.04 (50.33) | 13.80 (18.65) |

**Figure 4.**
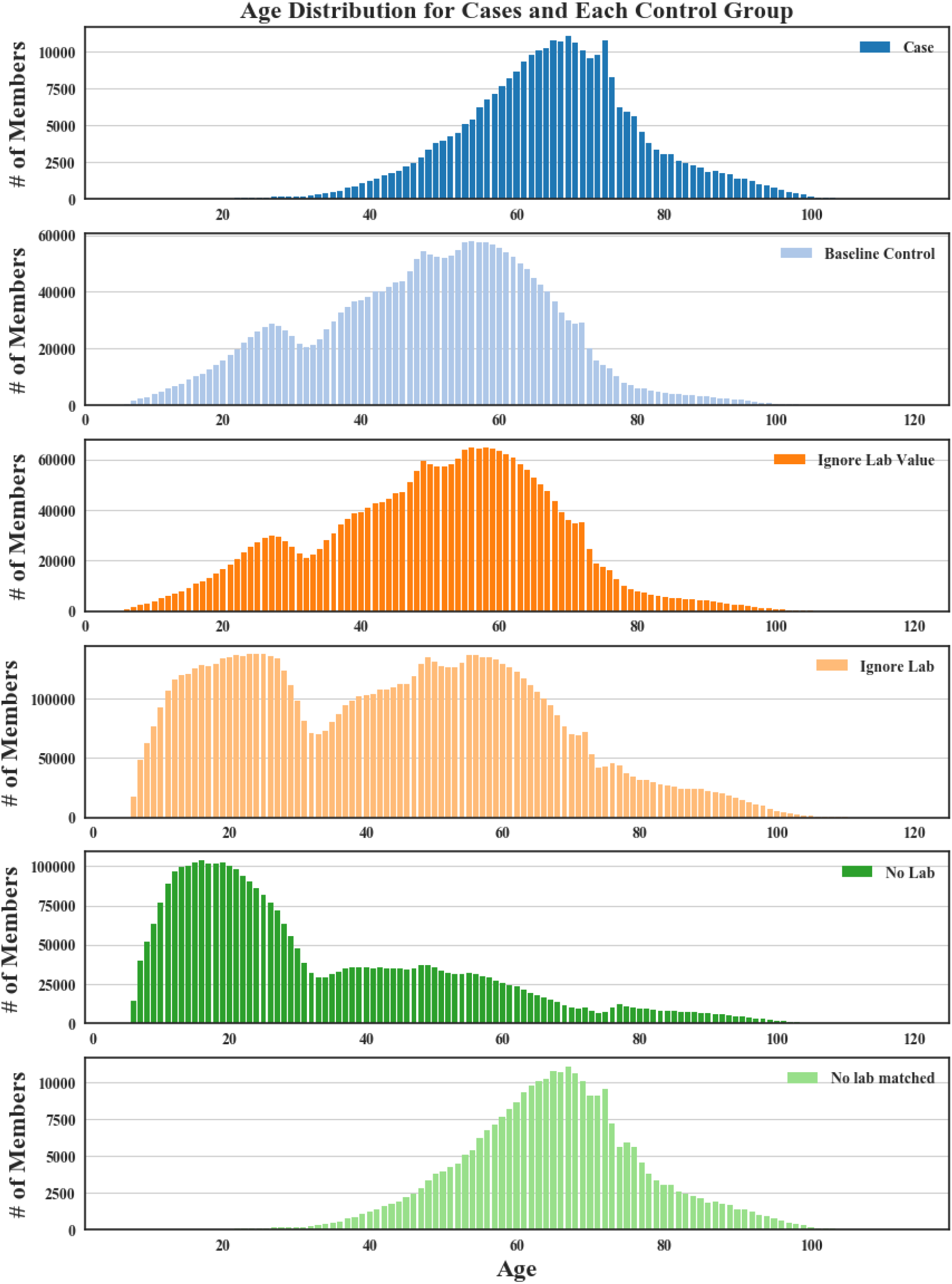
Age distributions under each different control definition.

We found a significant association between type 2 diabetes and depression in most bootstrapped samples. When sampling case/control cohorts at n=10,000, only 1 out of 1600 total randomizations resulted in a non-significant test. After matching, all 800 results in a significant finding. Within the matched populations, requiring that controls have a glucose test ordered but ignoring its value (median OR=1.291, 95% CI = [1.195, 1.397]) results in a slightly weaker association compared to the baseline control (median OR=1.248, 95% CI = [1.155, 1.349]). This reinforces the existence of a true association, as this variation may be inducing some case contamination in the control group. Testing against the no lab control group (median OR=2.792, 95% CI = [2.539, 3.069]) yields a much higher odds ratio compared to baseline. Testing against the ignore lab control group (median OR=1.361, 95% CI = [1.1257, 1.473]) results in an odds ratio between the baseline control and the no lab control group.

At lower sample sizes (n=2,000), the odds ratios estimates are more widely distributed (95% CI range is 2.23 times greater for the baseline group, 2.27 times greater for the ignore lab value group, 2.29 times greater for the ignore lab group, and 2.24 times greater for the no lab group). In the no lab control setting, reducing cohort size does not affect the significance of the test, given the margin between the OR and one. When the OR is closer to one, smaller cohort size and less precise estimate may lead to a change in direction for the OR and tests may lose significance. As many as 111/200 tests in the ignore lab value setting in Figure 5B tested non-significant.

**Figure 5.**
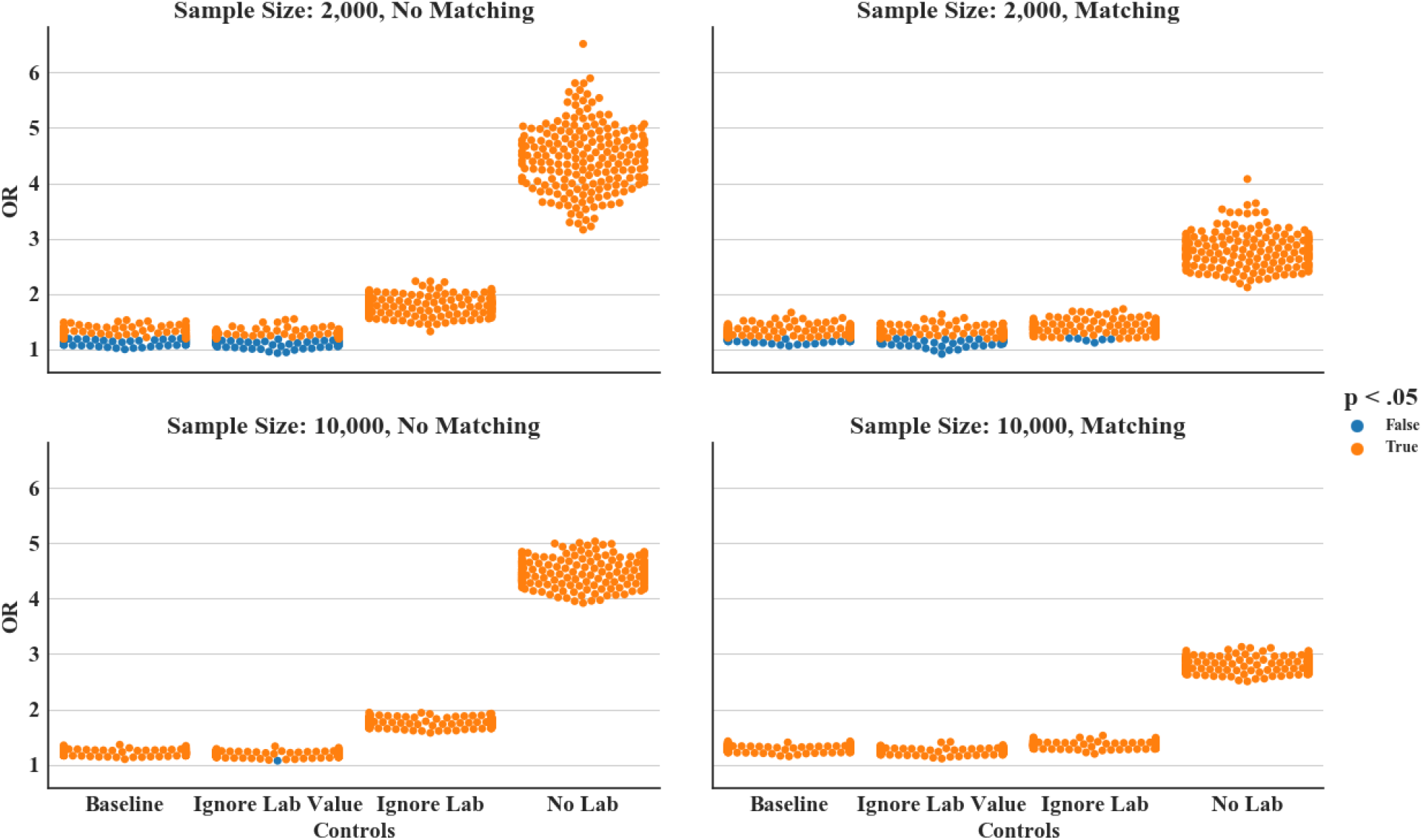
Odds ratio among each of the different control groups for 200 samples. **A.)** Sample size of 2,000 without matching, **B.)** Sample size of 2,000 with matching on age and sex, **C.)** Sample size of 10,000 without matching, **D.)** Sample size of 10,000 with matching on age and sex.

## Discussion

We applied retrospective case-control study design in a large administrative claims dataset to test for association between depression and T2D using multiple control group definitions. Taken altogether, the evidence suggests a true association exists between depression and T2D, but we do not yet attempt to determine directionality or causality. We found that permutations to the control definition which we believe to be reasonable led to changes in results. This was shown through shifts in the demographic structure of the control population, as well as differences in the OR of association. With age and sex matching, these differences are tempered, but still significant.

These results indicate that it would possible for bad actors to manipulate results based on RWE in a manner that is difficult to detect without knowing all experimental parameters used by the bad actors. It is therefore critical to establish best practices regarding transparency, neutrality, conflicts of interest, data provenance, pre-registration, cohort selection, sample sizes and reporting of results prior to using RWE as a major component of the regulatory process.

There were several limitations to this study. While we measure an association between depression and T2D, our study does not attempt to conclude whether depression elevates the risk of developing T2D or vice versa. Furthermore, it is unclear what biases are being encoded by the different definitions and how they drive changes in OR. Moreover, we made several approximations in repurposing the eMERGE T2D algorithm for claims data which may have affected its specificity. In addition to the lack of formal determination of family history, systematic biases may exist in the availability of lab test values. Finally, the use of a private insurance dataset selects against older segments of the population who may qualify for Medicare, as well as unemployed/lower socio-economic status segments who cannot afford private insurance.

Despite these limitations, this study demonstrates that effect size in retrospective association studies may be maximized by cherry-picking a control group. We suggest that the ability to strategically select a control group is not limited to association studies but extends to most RWE and retrospective studies. To mitigate the risk of publishing an unsound result, we recommend: 1.) RWE-based studies use and do not modify externally generated and validated pre-defined eligibility criteria, 2.) if not possible RWE-based studies should pre-register eligibility criteria and protocol prior to obtaining data, 3.) report all permutations tested with results for each permutation (potentially through the use of an independent audit system), 4.) avoid subsampling or report all subsamples with a variety of random seeds. In addition, when used as evidence for decisions with a financial impact, studies should be replicated by an independent neutral body.

